# StabilitySort: assessment of protein stability changes on a genome-wide scale to prioritise potentially pathogenic genetic variation

**DOI:** 10.1101/2021.11.28.470298

**Authors:** Aaron Chuah, Sean Li, Andrea Do, Matt A Field, T. Daniel Andrews

## Abstract

**Summary:** Missense mutations that change protein stability are strongly associated with human inherited genetic disease. With the recent availability of predicted structures for all human proteins generated using the AlphaFold2 prediction model, genome-wide assessment of the stability effects of genetic variation can, for the first time, be easily performed. This facilitates the interrogation of personal genetic variation for potentially pathogenic effects through the application of stability metrics. Here, we present a novel algorithm to prioritise variants predicted to strongly destabilise essential proteins, available as both a standalone software package and a web-based tool. We demonstrate the utility of this tool by showing that at values of the Stability Sort Z-score above 1.6, pathogenic, protein-destabilising variants from ClinVar are detected at a 58% enrichment, over and above the destabilising (but presumably non-pathogenic) variation already present in the HapMap NA12878 genome.

**Availability and Implementation:** StabilitySort is available as both a web service (http://130.56.244.113/StabilitySort/) and can be deployed as a standalone system (https://gitlab.com/baaron/StabilitySort).

## 1 Introduction

The ease with which individual genomes can be sequenced belies the complexities we presently face when interpreting this information (MacArthur et al., 2014; Rehm, 2017; Tarailo-Graovac et al., 2017; Whiffin et al., 2017). Currently, interpretation of personal genome data is clinically relevant for only a small proportion of the variants identified (Manolio et al., 2019), and for the remainder, we lack a deep understanding of the relationship between missense variation and potential functional effects. Despite its’ importance, to date, the interpretive power of instability effects of mutations on proteins has been limited by incomplete structural information covering the observed variation (only 16% of human proteins match a structure in the Protein Data Bank with greater than 95% identity (Porta-Pardo et al., 2021)). However, DeepMind’s AlphaFold 2 has generated structural predictions whose accuracy approaches those of experimentally determined structures (Jumper et al., 2021). This, accompanied with the release of near-exhaustive sets of protein models from human and other species (Tunyasuvunakool et al., 2021), means that it is now feasible to produce structure-based estimates of the stability effects for the vast bulk of genomic variation encountered in a personal genome or exome sequence. While it is not feasible to use the AlphaFold algorithm for *de novo* prediction of mutant protein structures, we are able to use predicted wild type structures as a template for assessing the potential structural consequences of mutations (Pak et al., 2021)(Akdel et al., 2021; Pak & Ivankov, 2021). StabilitySort uses the AlphaFold predictions of 3D structures as templates to conduct genome-wide assessment of protein stability changes due to genetic variation, in the absence of a similar, complete resource of experimentally obtained structures. While the AlphaFold dataset of 3D predictions are variable in quality (Thornton et al., 2021), the almost genomewide coverage of the resource (Porta-Pardo et al., 2021) presents an opportunity for routine interpretation of the protein stability effects of personal genome variation information.

## 2 Analysis Workflow

StabilitySort uses internal data from exhaustive pre-computation of the predicted ΔΔG of each possible human missense variant, genome-wide, using AlphaFold-predicted structures (Tunyasuvunakool et al., 2021) with Maestro (Laimer et al., 2015). Maestro was chosen based on benchmarking results and the additional feature that it can simultaneously predict stability due to multiple amino acid substitutions (Marabotti et al., 2021). StabilitySort initially calculates stability values due to single amino acid substitutions in a single protein, as in most individuals, almost all proteins will harbour at most only a single variant. However, the user is also provided the option to pool all missense variation across a protein, should there be multiple variants in a given protein, to re-compute the predicted change in stability due to multiple substitutions.

The StabilitySort methodology seeks to identify missense variants that introduce either highly destabilising or stabilising changes into the encoded protein. We measure this effect by viewing the change with reference to the missense variation in the same protein from the GnomAD database (Karczewski et al., 2020). For each missense variant observed from an input genome, a StabilitySort Z-score test statistic is calculated that assesses the predicted ΔΔG with respect to all other non-disease associated variation identified in this protein. The StabilitySort Z-score measures how unusual a particular amino acid substitution is, in a particular protein, compared to the distribution of observed GnomAD substitutions in that same protein. Higher Z-score values indicate a higher chance that the amino acid substitution has a functional impact on the protein.

## 3 Results

To demonstrate the effectiveness of StabilitySort in prioritising disease-causing missense variants, we compared 1028 randomly sampled ClinVar pathogenic variants (annotated by ClinVar as CLNSIG=Pathogenic; (Landrum et al., 2020)) with the missense variation present in HapMap individual NA12878 (from the International Genome Sample Resource (Fairley et al., 2020)). The StabilitySort Z-score metric identified an enrichment of a subset of ClinVar variants that were unusually destabilising, given the population variation in these proteins (Supplementary Figure 1a). At Z-scores values greater than 1.6 there are 58% more ClinVar pathogenic mutations than at the same Z-score cutoff in the NA12878 genome. This enrichment was asymmetric and was only observed for destabiling amino acid substitutions. Furthermore, the predicted ΔΔG values alone did not show an excess of destabilising amino acid substitutions for the ClinVar pathogenic variants compared to the NA12878 variation (Supplementary Figure 1b). Interpretation of predicted ΔΔG in the context of gene importance is informative, as the range of stability effects observed increases as proteins become more tolerant to loss-of-function mutations (redundant and/or non-essential genes), though the median ΔΔG does not significantly vary (Supplementary Figure 1c; see notches in bar-plot). StabilitySort did not identify candidate pathogenic variation with strong stability effects in the genome of NA12878.

## 4 Conclusion

We present StabilitySort, a genetic variation prioritisation tool for the genome-wide detection of protein stability effects that may contribute to disease. This system is available as both a web service and as standalone software. With this methodology, it is now possible to scan for the presence of likely pathogenic stabilising or destabilising protein mutations in a high-throughput manner.

## Figures

**Figure 1.**
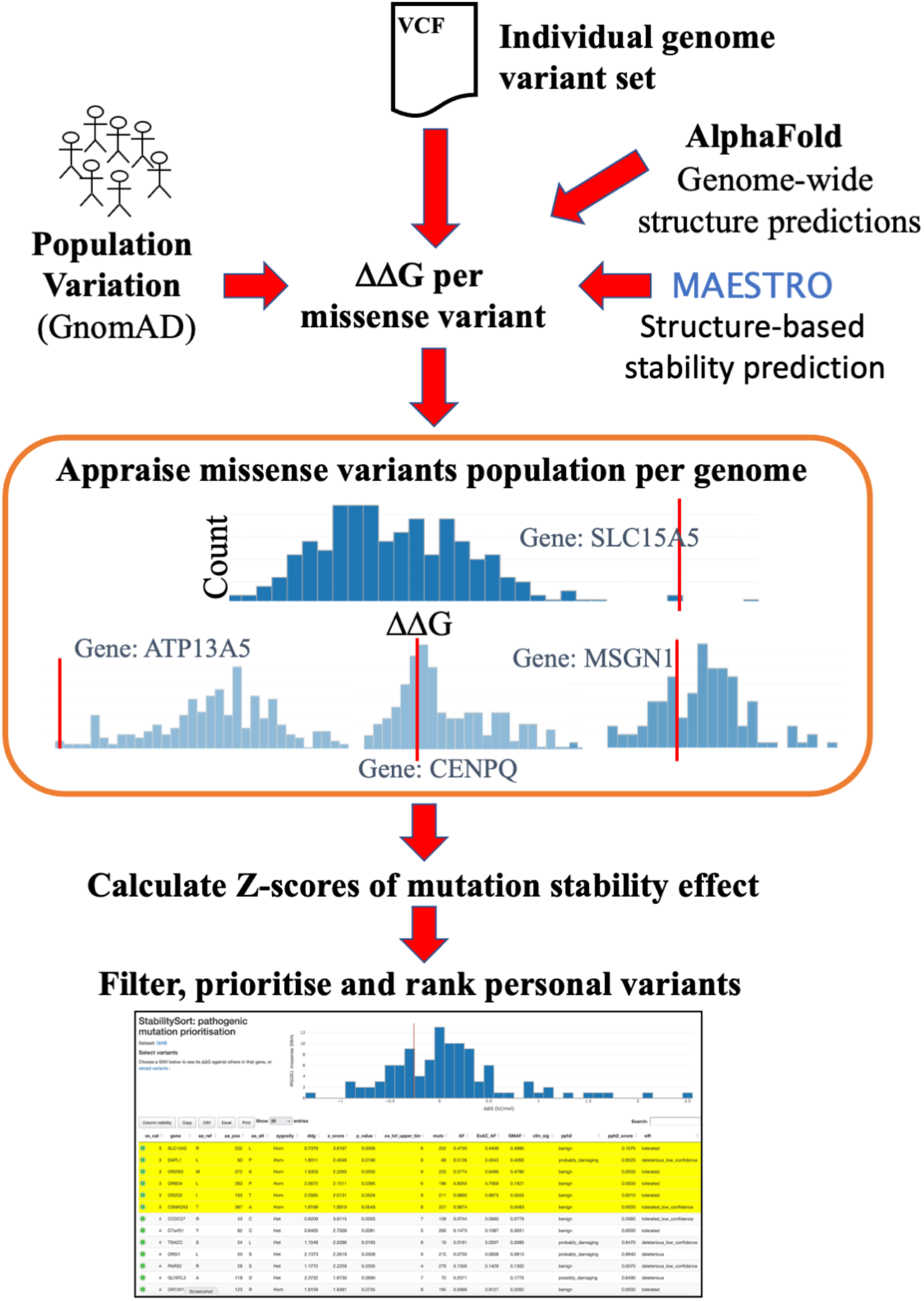
StabilitySort allows prioritisation of the full missense variant set of an given genome through an automated workflow that annotates amino acid subsitutions with their predicted stability effects. Input to the workflow begins with a user-supplied VCF file of variation. An index of exhaustively predicted ΔΔG values, predicted with the Maestro algorithm, using AlphaFold2-predicted human protein structures, is used to compare the value of the variant in question with the variation observed in the protein described in the GnomAD database. This comparison is quantified with a Z-score, and this metric, along with other annotated values, can be used to prioritise and order potentially-pathogenic missense variation at a genome-wide scale.

**Supplementary Figure 1.**
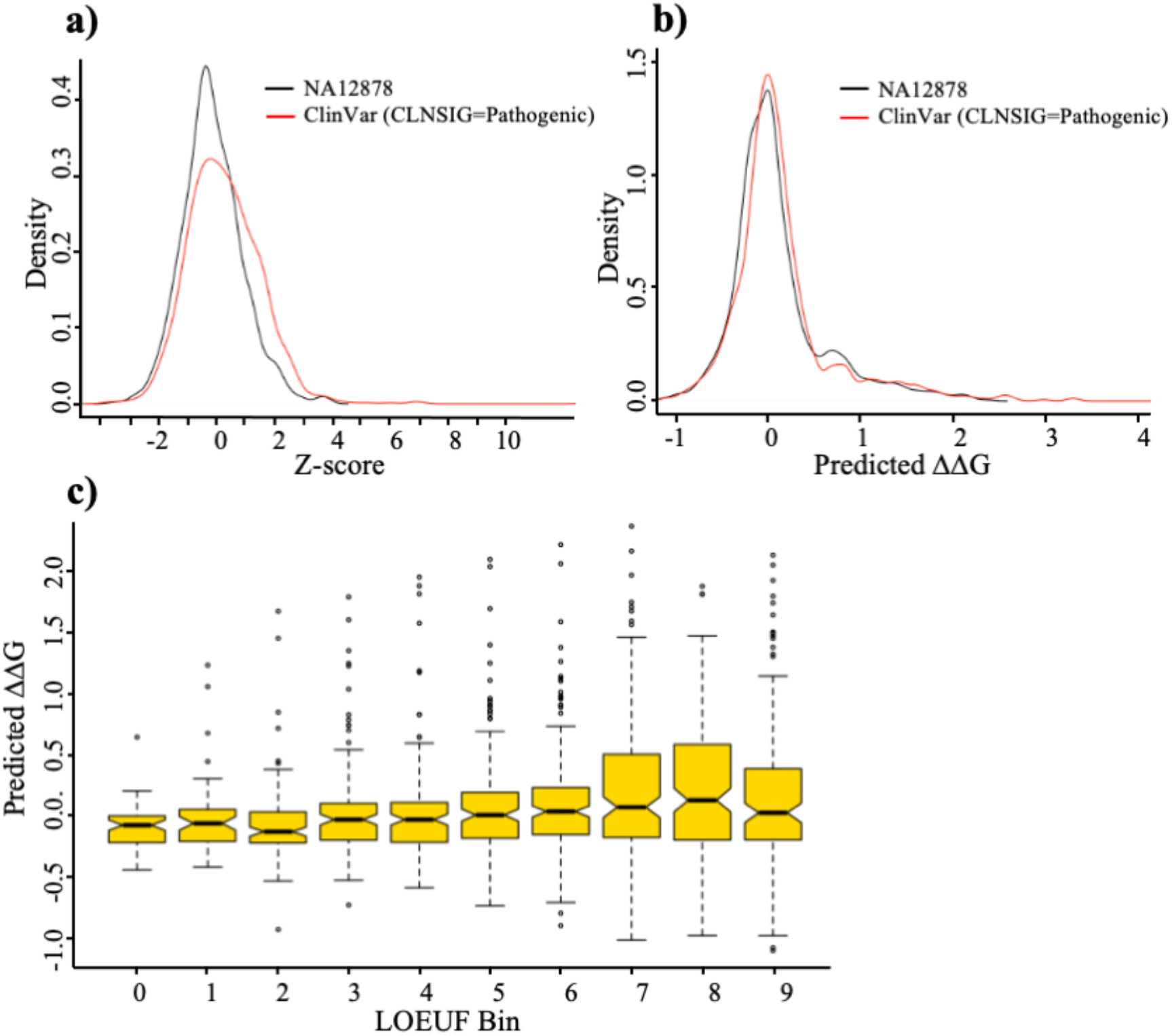
Comparison of ClinVar pathogenic missense variants with the personal missense variation identified in the NA12878 HapMap genome. **a)** Comparison of distribution of change of stability Z-Scores, and **b)** MAESTRO predicted ΔΔG values between missense variant sets. **c)** Distribtions of predicted ddG values for all missense variants from the NA12878 genome, separated into bins by LOUEF decile values. Overlap of notches in bar-plot indicate similarity of means between bins in a 95% confidence interval. For each bar coloured in gold, the bold mid-line indicates the median predicted DDG, the upper and lower ends of the bar indicate the second and third quartile and the whiskers indicate the first and fourth quartiles. The oulier values are indicated with dots. The ten bins separated by LOUEF decile values are numbered (0-9) according to increasing tolerance of loss-of-function mutations (see (Karczewski et al., 2020)). Bin 0 contains the least tolerant of loss-of-function mutations (designated as essential genes) and bin 9 contains the most tolerant (designated redundant or non-essential genes).

## References

Akdel, M., Pires, D. E. V, Pardo, E. P., Jänes, J., Zalevsky, A. O., Mészáros, B., Bryant, P., Good, L. L., Laskowski, R. A., Pozzati, G., Shenoy, A., Zhu, W., Kundrotas, P., Serra, V. R., Rodrigues, C. H. M., Dunham, A. S., Burke, D., Borkakoti, N., Velankar, S., … Beltrao, P. (2021). A structural biology community assessment of AlphaFold 2 applications. BioRxiv, 2021.09.26.461876. https://doi.org/10.1101/2021.09.26.461876

Fairley, S., Lowy-Gallego, E., Perry, E., & Flicek, P. (2020). The International Genome Sample Resource (IGSR) collection of open human genomic variation resources. Nucleic Acids Research, 48(D1), D941–D947. https://doi.org/10.1093/NAR/GKZ836

Jumper, J., Evans, R., Pritzel, A., Green, T., Figurnov, M., Ronneberger, O., Tunyasuvunakool, K., Bates, R., Žídek, A., Potapenko, A., Bridgland, A., Meyer, C., Kohl, S. A. A., Ballard, A. J., Cowie, A., Romera-Paredes, B., Nikolov, S., Jain, R., Adler, J., … Hassabis, D. (2021). Highly accurate protein structure prediction with AlphaFold. Nature 2021, 1–11. https://doi.org/10.1038/s41586-021-03819-2

Karczewski, K. J., Francioli, L. C., Tiao, G., Cummings, B. B., Alföldi, J., Wang, Q., Collins, R. L., Laricchia, K. M., Ganna, A., Birnbaum, D. P., Gauthier, L. D., Brand, H., Solomonson, M., Watts, N. A., Rhodes, D., Singer-Berk, M., England, E. M., Seaby, E. G., Kosmicki, J. A., … MacArthur, D. G. (2020). The mutational constraint spectrum quantified from variation in 141,456 humans. Nature 2020 581:7809, 581(7809), 434–443. https://doi.org/10.1038/s41586-020-2308-7

Laimer, J., Hofer, H., Fritz, M., Wegenkittl, S., & Lackner, P. (2015). MAESTRO - multi agent stability prediction upon point mutations. BMC Bioinformatics, 16(1), 1–13. https://doi.org/10.1186/S12859-015-0548-6/TABLES/5

Landrum, M. J., Chitipiralla, S., Brown, G. R., Chen, C., Gu, B., Hart, J., Hoffman, D., Jang, W., Kaur, K., Liu, C., Lyoshin, V., Maddipatla, Z., Maiti, R., Mitchell, J., O’Leary, N., Riley, G. R., Shi, W., Zhou, G., Schneider, V., … Kattman, B. L. (2020). ClinVar: improvements to accessing data. Nucleic Acids Research, 48(D1), D835–D844. https://doi.org/10.1093/NAR/GKZ972

MacArthur, D. G., Manolio, T. A., Dimmock, D. P., Rehm, H. L., Shendure, J., Abecasis, G. R., Adams, D. R., Altman, R. B., Antonarakis, S. E., Ashley, E. A., Barrett, J. C., Biesecker, L. G., Conrad, D. F., Cooper, G. M., Cox, N. J., Daly, M. J., Gerstein, M. B., Goldstein, D. B., Hirschhorn, J. N., … Gunter, C. (2014). Guidelines for investigating causality of sequence variants in human disease. In Nature (Vol. 508, Issue 7497, pp. 469–476). https://doi.org/10.1038/nature13127

Manolio, T. A., Rowley, R., Williams, M. S., Roden, D., Ginsburg, G. S., Bult, C., Chisholm, R. L., Deverka, P. A., McLeod, H. L., Mensah, G. A., Relling, M. V., Rodriguez, L. L., Tamburro, C., & Green, E. D. (2019). Opportunities, resources, and techniques for implementing genomics in clinical care. The Lancet, 394(10197), 511–520. https://doi.org/10.1016/S0140-6736(19)31140-7

Marabotti, A., Scafuri, B., & Facchiano, A. (2021). Predicting the stability of mutant proteins by computational approaches: an overview. Briefings in Bioinformatics, 22(3). https://doi.org/10.1093/BIB/BBAA074

Pak, M. A., & Ivankov, D. N. (2021). Best templates outperform homology models in predicting the impact of mutations on protein stability. BioRxiv, 2021.08.26.457758. https://doi.org/10.1101/2021.08.26.457758

Pak, M. A., Markhieva, K. A., Novikova, M. S., Petrov, D. S., Vorobyev, I. S., Maksimova, E. S., Kondrashov, F. A., & Ivankov, D. N. (2021). Using AlphaFold to predict the impact of single mutations on protein stability and function. BioRxiv, 2021.09.19.460937. https://doi.org/10.1101/2021.09.19.460937

Porta-Pardo, E., Ruiz-Serra, V., & Valencia, A. (2021). The structural coverage of the human proteome before and after AlphaFold. BioRxiv, 2021.08.03.454980. https://doi.org/10.1101/2021.08.03.454980

Rehm, H. L. (2017). A new era in the interpretation of human genomic variation. In Genetics in Medicine. https://doi.org/10.1038/gim.2017.90

Tarailo-Graovac, M., Zhu, J. Y. A., Matthews, A., van Karnebeek, C. D. M., & Wasserman, W. W. (2017). Assessment of the ExAC data set for the presence of individuals with pathogenic genotypes implicated in severe Mendelian pediatric disorders. Genetics in Medicine 2017 19:12, 19(12), 1300–1308. https://doi.org/10.1038/gim.2017.50

Thornton, J. M., Laskowski, R. A., & Borkakoti, N. (2021). AlphaFold heralds a data-driven revolution in biology and medicine. Nature Medicine 2021 27:10, 27(10), 1666–1669. https://doi.org/10.1038/s41591-021-01533-0

Tunyasuvunakool, K., Adler, J., Wu, Z., Green, T., Zielinski, M., Žídek, A., Bridgland, A., Cowie, A., Meyer, C., Laydon, A., Velankar, S., Kleywegt, G. J., Bateman, A., Evans, R., Pritzel, A., Figurnov, M., Ronneberger, O., Bates, R., Kohl, S. A. A., … Hassabis, D. (2021). Highly accurate protein structure prediction for the human proteome. Nature 2021 596:7873, 596(7873), 590–596. https://doi.org/10.1038/s41586-021-03828-1

Whiffin, N., Minikel, E., Walsh, R., O’Donnell-Luria, A. H., Karczewski, K., Ing, A. Y., Barton, P. J. R., Funke, B., Cook, S. A., Macarthur, D., & Ware, J. S. (2017). Using high-resolution variant frequencies to empower clinical genome interpretation. Genetics in Medicine 2017 19:10, 19(10), 1151–1158. https://doi.org/10.1038/gim.2017.26

